# Effects of TP63 Mutations on Keratinocyte Adhesion and Migration

**DOI:** 10.1101/2023.05.04.539104

**Authors:** Maddison N. Salois, Jessica A. Gugger, Saiphone Webb, Christina E. Sheldon, Shirley P. Parraga, G. Michael Lewitt, Dorothy K. Grange, Peter J. Koch, Maranke I. Koster

## Abstract

The goal of this study was to investigate the molecular mechanisms responsible for the formation of skin erosions in patients affected by Ankyloblepharon-ectodermal defects-cleft lip/palate syndrome (AEC). This ectodermal dysplasia is caused by mutations in the *TP63* gene, which encodes several transcription factors that control epidermal development and homeostasis. We generated induced pluripotent stem cells (iPSC) from AEC patients and corrected the *TP63* mutations using genome editing tools. Three pairs of the resulting conisogenic iPSC lines were differentiated into keratinocytes (iPSC-K). We identified a significant downregulation of key components of hemidesmosomes and focal adhesions in AEC iPSC-K compared to their gene-corrected counterparts. Further, we demonstrated reduced iPSC-K migration, suggesting the possibility that a process critical for cutaneous wound healing might be impaired in AEC patients. Next, we generated chimeric mice expressing a *TP63-AEC* transgene and confirmed a downregulation of these genes in transgene-expressing cells *in vivo*. Finally, we also observed these abnormalities in AEC patient skin. Our findings suggest that integrin defects in AEC patients might weaken the adhesion of keratinocytes to the basement membrane. We propose that reduced expression of extracellular matrix adhesion receptors, potentially in conjunction with previously identified desmosomal protein defects, contribute to skin erosions in AEC.

## BACKGROUND

Ankyloblepharon-ectodermal defects-cleft lip/palate syndrome (AEC; OMIM #106260) (Figure 1A) is an ectodermal dysplasia characterized by skin fragility leading to the formation of severe skin erosions ^1^. AEC is most frequently caused by missense mutations in the *TP63* gene, which encodes several transcription factors with essential roles in epidermal and appendage development ^1-3^. Currently, only symptomatic treatment is available for the devastating skin erosions. We and others have previously identified abnormalities in keratinocyte differentiation, cell-cell adhesion, and cell-extracellular matrix (ECM) adhesion in small cohorts of AEC patients ^4-9^. To facilitate the design of future therapeutic options, it is essential to gain a better understanding of the pathophysiological mechanisms that contribute to skin fragility in this disorder. We have previously developed a human stem cell-based model system to study AEC keratinocytes ^6^. We generated induced pluripotent stem cells (iPSC) from three AEC patients using an integration-free viral reprogramming system ^6,10^. Each of these iPSC lines harbors a single point mutation in exons 13 or 14 of *TP63* (F513S, I537T, R598L), which encode the SAM domain ^11^. This domain is believed to be required for TP63 interactions with transcriptional coregulators, although very few interacting proteins have been identified thus far ^12-14^. To repair the *TP63* mutations in patient-derived iPSC, we used Crispr/Cas-mediated and TALEN-mediated genome editing, thereby establishing gene-corrected (GC) iPSC lines ^6,15^. We used a well-established protocol that relies on BMP4 and retinoic acid to differentiate iPSC into iPSC-derived keratinocytes (iPSC-K) ^6,16^. We previously demonstrated that these iPSC-K express marker proteins of normal human keratinocytes ^6,16^. Further, using these AEC and GC iPSC-K, we identified major defects in the expression and subcellular distribution of desmosomal proteins ^6^. These defects were also observed in AEC patient skin, further validating AEC iPSC-K as a model for AEC skin. Here, we identify additional defects in the expression and the function of hemidesmosomal and focal adhesion proteins in AEC iPSC-K and in AEC patient skin. Loss of expression or autoimmune targeting of many of the affected genes, such as *ITGA6, ITGB4* and *COL17A1*, have previously been linked to severe subepidermal blistering diseases ^17,18^. Consequently, our findings suggest that defects in basement membrane anchorage might contribute to skin fragility in AEC. Further, many of the integrins affected in AEC have previously also been linked to signaling pathways that control keratinocyte fate and function. It is thus likely that the deregulated expression of integrins in AEC might affect a wide range of keratinocyte properties, such as migration and differentiation.

**Figure 1:**
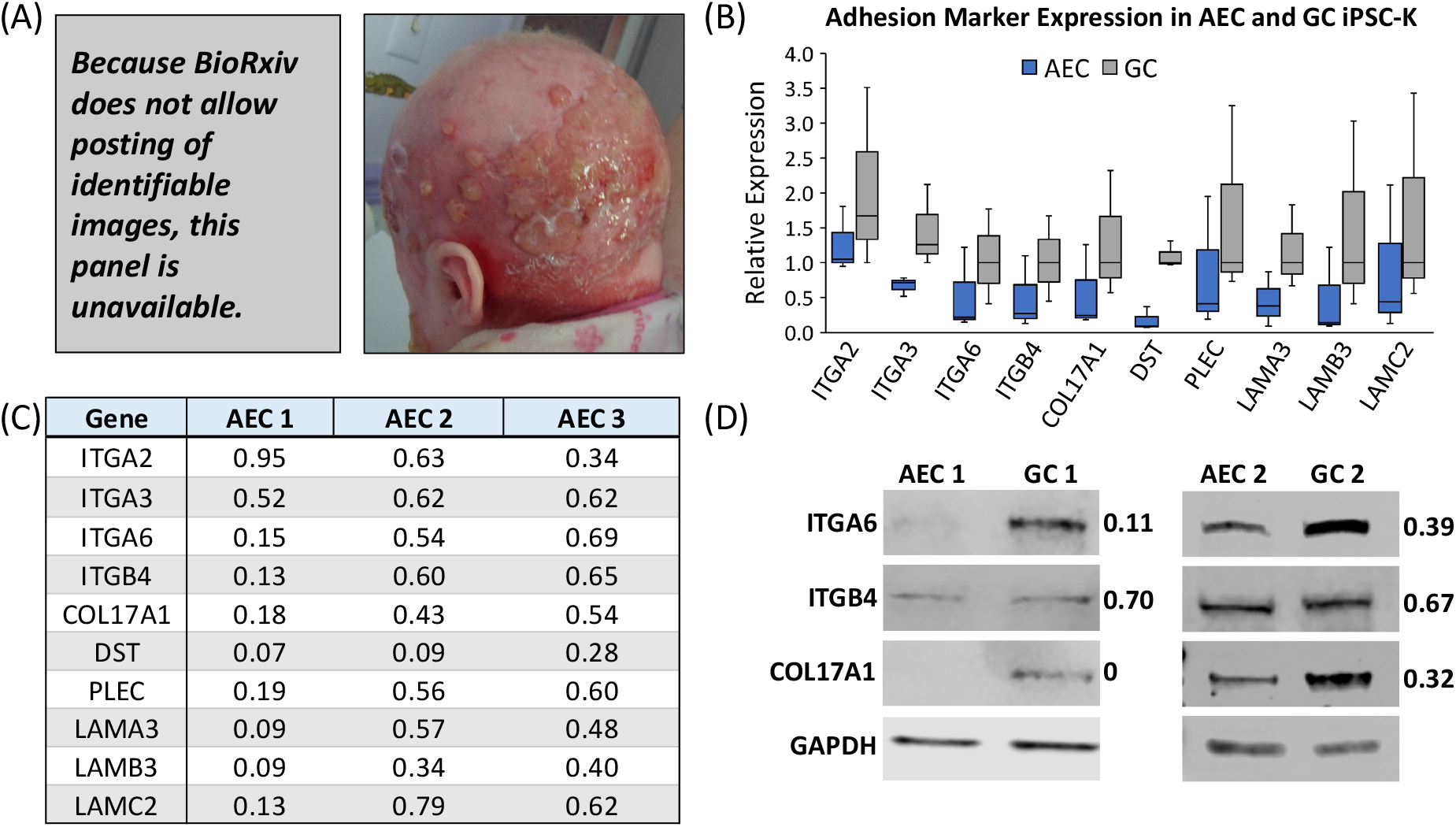
Hemidesmosomal and focal adhesion components are downregulated in AEC iPSC-K. (A) Images of an AEC patient showing skin erosions. Note the superficial erosions, abnormal and excessive granulation tissue and residual scarring alopecia (left) and the severe scalp erosive dermatitis (with overlying bacterial infection) and patchy alopecia with light wiry hairs (right). The photos are a courtesy of the National Foundation for Ectodermal Dysplasias (NFED) and are shown with parental consent. (B, C) qRT-PCR analysis of three pairs of AEC and GC iPSC-K. (B) All gene expression values were normalized to the same GC iPSC-K sample (GC1). The horizontal line represents the median; the whiskers represent the highest and lowest values in each dataset. (C) Numerical values of qRT-PCR analysis. In contrast to (B), all AEC iPSC-K gene expression values were normalized to their respective GC iPSC-K. Note the downregulation of all genes tested in (B) and (C). (D) Example of Western blot analysis using two pairs of AEC and GC iPSC-K. Numbers indicate ratios of signal intensity of AEC iPSC-K to conisogenic GC iPSC-K. The observed variability in transcript and protein expression between the three GC iPSC-K lines is expected, and reflects the expression variability of cell surface receptors often observed between different individuals ^26,27^.

## QUESTIONS ADDRESSED

The goal of this investigation was to identify mechanisms contributing to the skin fragility observed in AEC patients.

## EXPERIMENTAL DESIGN

See Supplemental Information for experimental procedures.

## RESULTS

We used three conisogenic pairs of iPSC-derived patient (AEC iPSC-K) and gene-corrected (GC iPSC-K) keratinocytes (*TP63* mutations: F513S, I537T, R598L). These iPSC-K were previously generated by our laboratories, and all research participants donating skin biopsies provided their written informed consent/assent prior to their inclusion in the study ^6^. The iPSC-K were cultured under conditions that maintain keratinocytes in a proliferative, undifferentiated state to mimic basal keratinocytes of the epidermis ^19,20^. We previously established that iPSC-K generated using our method have similar characteristics to normal human keratinocytes as determined by transcriptome analyses, protein expression, and their potential to differentiate into keratinocytes representing the different layers of the epidermis ^6,16,21^. iPSC-K cultures were considered appropriate for analysis if at least 95% of the cells expressed basal keratinocyte markers, such as KRT14 and TP63 ^16^. We next conducted RNA sequencing of the three conisogenic iPSC-K pairs, referred to here as AEC1-3 and GC1-3. Using Gene Set Enrichment Analysis (GSEA) and Genome Ontology (GO) analysis ^22,23^, we found that two of the most significantly affected groups of genes were classified as “integrin family cell surface interactions” and “extracellular matrix”. Specifically, we observed a downregulation of several genes associated with the interaction of basal keratinocytes with the extracellular matrix (ECM). Expression levels of the hemidesmosomal transmembrane receptor genes *ITGA6, ITGB4* and *COL17A1* as well as genes encoding the cytoplasmic hemidesmosomal adaptor proteins *DST* and *PLEC* were all reduced in AEC iPSC-K. In addition, genes encoding the hemidesmosomal ligands *LAMA3, LAMB3* and *LAMC2* were also downregulated. These genes encode proteins that are essential for keratinocyte adhesion and migration ^24,25^. Mutations in these genes or auto-antibodies generated against the encoded proteins are responsible for blistering skin disorders, further underscoring their importance ^17,18^. As shown in Figures 1B-D, we independently validated the RNAseq data showing downregulation of important cell-ECM adhesion genes and proteins in AEC iPSC-K using qRT-PCR and Western Blot analysis. These results suggest that TP63 is an important regulator of the hemidesmosomal adhesion complex. Further, the focal adhesion integrins ITGA2 and ITGA3 were reduced in their expression, suggesting that adhesion, migration, and integrin-mediated signaling processes might be perturbed in AEC keratinocytes. Importantly, all genes were downregulated in each AEC iPSC-K line compared to their respective GC counterparts (Figure 1C). The expression variability among different GC iPSC-K lines was expected, as it has previously been shown that cell surface receptors and their ligands show high variability in their expression levels between individuals ^26,27^.

To validate our in vitro observations, we assessed the expression of integrins in chimeric mice expressing a *TP63-AEC* transgene carrying the AEC3 mutation (Figure 2A; I537T mutation). These mice were generated using mouse embryonic stem (ES) cells transduced with a lentivirus encoding the *TP63-AEC* transgene. The *TP63-AEC* transgene was placed under the control of a CAG promoter and was linked to a TdTomato reporter gene via a T2A domain, a self-cleaving peptide that allows for the equimolar expression of the *TP63-AEC* transgene and the TdTomato reporter (Figure 2A) ^28^. Transduced ES cells were isolated using Fluorescence Activated Cell Sorting (FACS) using TdTomato expression as a surrogate marker for the presence of the *TP63-AEC* transgene. Transduced and FACS-sorted ES cells were then injected into mouse blastocysts followed by implantation into pseudo-pregnant recipient mice. Chimeric embryos were analyzed at embryonic day (E) 18.5. This developmental stage was chosen to avoid potential perinatal lethality caused by the transgene. A severe AEC phenotype could result in cleft lip/palate development, which is predicted to be lethal in mice due to an inability to nurse ^29,30^. The TdTomato marker was used to identify cells expressing the *TP63-AEC* transgene within the chimeric mouse tissue via immunofluorescence microscopy, enabling us to compare the properties of mutant and wild type cells side by side *in vivo*. We observed a reduction in integrin expression in cells expressing the *TP63-AEC* transgene, in several tissues, including the epidermis and the palate (Figure 2B-E). Our analysis in mouse and human (below) tissues was somewhat limited due to the unavailability of antibodies that stain paraffin-preserved tissues. This chimeric animal model will be an essential tool to understand how *TP63-AEC* mutations affect development of the epidermis, skin appendages, and other TP63-expressing epithelial tissues while avoiding potentially lethal effects of the *TP63* mutations in mice. Further, our approach can easily be expanded in future studies to compare the molecular pathology of different AEC mutations *in vivo* in a time- and cost-effective manner.

**Figure 2:**
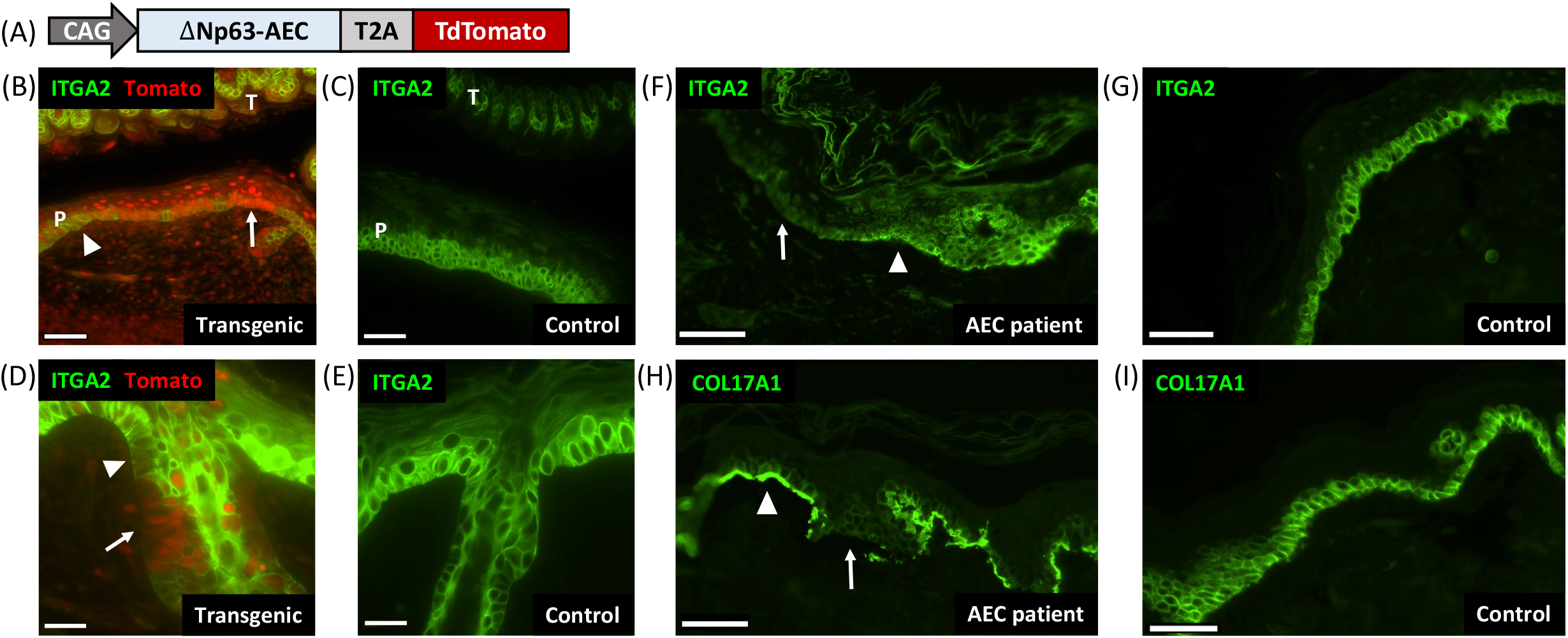
Downregulation of cell surface receptors in mutant TP63-expressing mouse tissues and AEC patient skin. (A) Schematic representation of the lentiviral construct used to generate chimeric mice expressing mutant *TP63*. The construct is designed to express mutant TP63 and TdTomato in equimolar amounts due to the presence of the T2A self-cleaving peptide. The CAG promoter is active in embryonic stem cells as well as epidermal and dermal skin cells. Transgenic cells can be identified in tissues by TdTomato expression. (B-E) Immunofluorescent staining of mouse tissue with antibodies against TdTomato (red) and ITGA2 (green). (B) Chimeric mouse palate (P) and tongue (T) are shown. Note the downregulation of ITGA2 (green) in areas of the palate that express the transgene (red; arrow). Arrowheads indicate non-transgenic areas with normal ITGA2 expression. (C) Control mouse palate and tongue. (D) Chimeric mouse skin. Note the downregulation of ITGA2 (green) in the transgenic (red) upper permanent portion of the hair follicle (arrow). (E) Control mouse skin. Size bars in (B, C) 50 μm, size bars in (D, E) 20 μm. (F, G) Immunofluorescent staining for ITGA2 on (F) AEC patient skin and (G) control skin. (H, I) Immunofluorescent staining for COL17A1 on (H) AEC patient skin and (I) control skin. As above, arrowheads indicate normal expression; arrows indicate loss of expression. Note the focal loss of ITGA2 and COL17A1 in AEC patient skin. Size bars in (F-I) 50 μm.

Next, we assessed basal keratinocyte protein expression in AEC patient skin. We observed a focal downregulation of ITGA2 and COL17A1 in AEC patient skin (Figures 2F-I). Note that the mechanistic basis for the focal downregulation apparent in Figures 2F and H is currently unknown. Interestingly, focal loss of desmosomal proteins and of suprabasal keratins expressed in terminally differentiating keratinocytes has been observed in AEC patient skin as well ^5,6,8^. Compensatory mechanisms must exist *in vivo* to restore or maintain expression of TP63 target genes in focal cell clusters of AEC patient epidermis. Identifying these mechanisms might lead to new approaches in treating this devastating disease.

Integrins have critical roles in cell-ECM adhesion and in cell migration ^31^. As specific integrin pairs have different ECM ligand binding capabilities ^31,32^ (Figure 3A), we next assessed the ability of AEC iPSC-K to adhere to and migrate on different ECM substrates. To this end, we coated tissue culture plates with one of three ECM substrates: LAM332, LAM511, or collagen IV (COL4). As shown in Figure 3A, these matrix proteins are ligands of the integrins that are downregulated in AEC iPSC-K. For the adhesion assays, fluorescently labeled AEC and GC iPSC-K were seeded on these plates and allowed to adhere for 30 minutes. We then calculated the percentage of cells that had adhered to the ECM-coated well. As shown in Figure 3B, AEC iPSC-K showed a reduced ability to adhere to all three ECM substrates, a finding that is consistent with the reduced expression of cell surface integrin receptors in AEC iPSC-K. Similarly, we found that AEC iPSC-K showed reduced migration on all ECM substrates ^33^ (Figures 3C,D). Collectively, these results demonstrate that AEC iPSC-K show reduced adhesion to, and migration on, various ECM substrates, a pathological process most likely caused by the reduced cell surface expression of several integrins.

**Figure 3:**
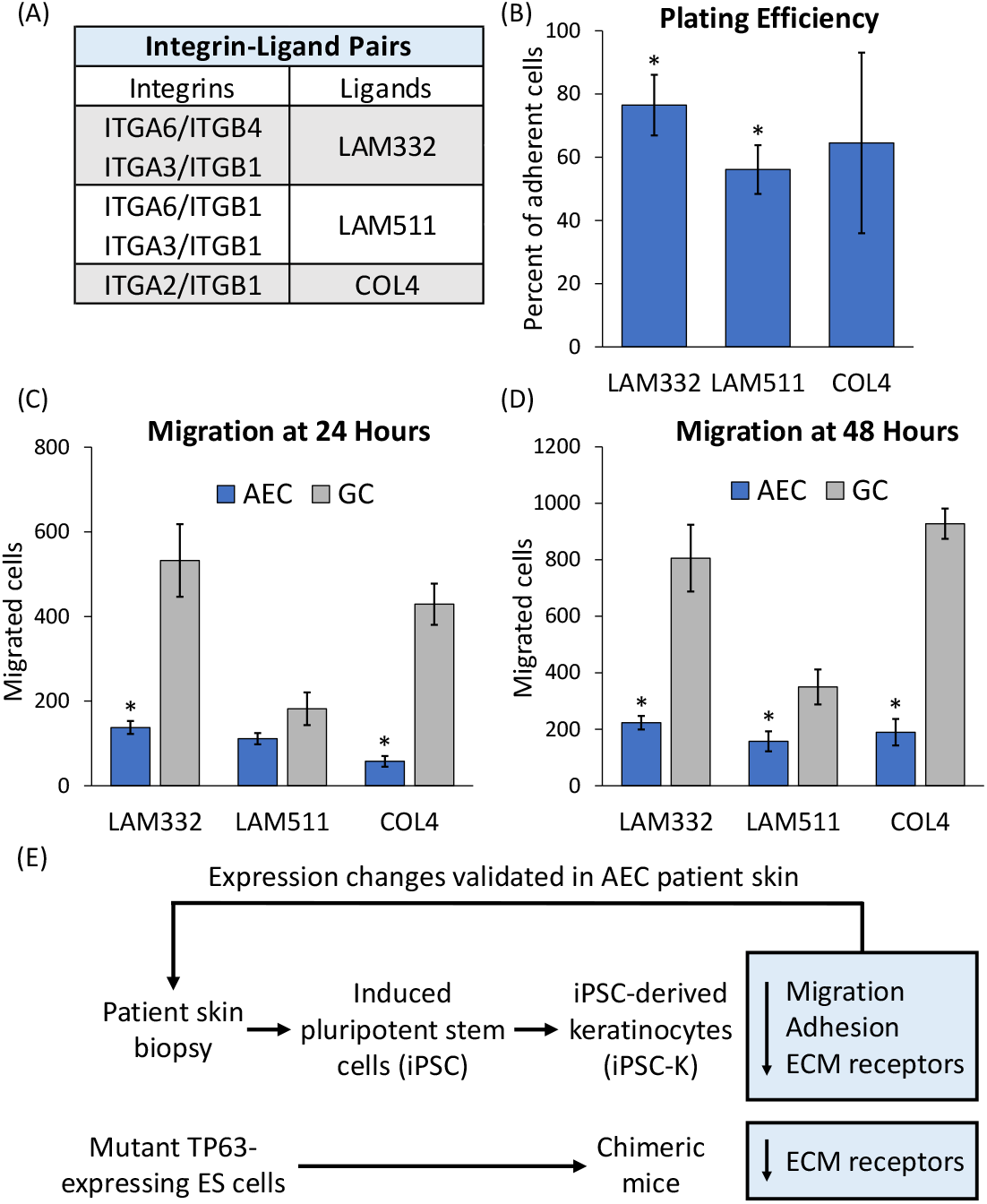
Reduced plating efficiency and migration of AEC iPSC-K. (A) Table indicating integrins and their ligands in the epidermis. (B) Plating efficiency was determined by calculating the relative percentage of AEC iPSC-K versus GC iPSC-K that adhered to the indicated ECM substrates. Plating efficiency of GC iPSC-K was set to 100%. (C, D) Assessment of AEC and GC iPSC-K migration on different ECM substrates at (C) 24 and (D) 48 hours. Note that both plating efficiency and migration were reduced in AEC iPSC-K on all tested substrates. ^*^p<0.05. (E) Schematic representation of the outline of our experimental approach and the main findings of this study.

## CONCLUSIONS AND PERSPECTIVES

In the present study, we demonstrate that keratinocytes carrying *TP63-AEC* mutations show a coordinated downregulation of hemidesmosomal proteins and focal adhesion proteins (Figure 3E). Further, we demonstrate that several of these proteins are also downregulated in an animal model for AEC as well as in AEC patient skin (Figure 3E). In addition to the observed defects in cell-ECM adhesion, we and others have previously identified abnormalities in cell-cell adhesion (desmosomal abnormalities) in AEC-derived keratinocytes and in AEC patient skin ^6,7^. Interestingly, mutations in cell adhesion genes or auto-antibodies generated against the encoded proteins are responsible for several severe blistering skin disorders ^17,18,34^. Thus, our data suggest that the collective dysfunction of multiple adhesion systems is responsible for skin erosions in AEC patients.

It is noteworthy that several of the hemidesmosomal and focal adhesion proteins are simultaneously downregulated in AEC-derived keratinocytes. These data suggest that TP63 is a key regulator of these genes. This is supported by the finding that TP63 interacts with the promoters and enhancers of many keratinocyte adhesion genes in human keratinocytes, including the downregulated genes we identified in this study ^7,35,36^. At this time, we do not know which of the downregulated genes are the major drivers of the skin fragility phenotype. Further experiments will be aimed at determining whether restoration of specific adhesion proteins can restore adhesion, migration, and differentiation properties in AEC-derived keratinocytes.

Finally, the focal nature of adhesion protein downregulation in AEC patient skin suggests the existence of mechanisms that restore or maintain hemidesmosomal and focal adhesion proteins in the presence of *TP63* mutations. There is currently no evidence for the existence of a genetic mechanism, such as revertant mosaicism ^37^, to explain this phenomenon. Regardless of the genetic or epigenetic mechanism, the ability to restore adhesion is consistent with the observation that AEC skin erosions heal over time and are generally small or absent in adult patients. Identifying the mechanisms responsible for the restoration of skin integrity might lead to the development of novel therapeutic approaches.

## Ethical statements

All animal experiments were approved by the Institutional Animal Care and Use Committee (IACUC) of East Carolina University under AUP #208 (approval date 5/6/2020). Skin biopsies from AEC patients and control individuals were obtained under IRB protocols #17240 through St Louis University (approval date 6/7/2011), #19-0501 through the University of Colorado Denver (approval date 4/19/2019), and #201901088 through Washington University (approval date 5/13/2019). All research participants provided informed written consent. The patient photographs shown were used with written consent of the patient’s parents.

## Acknowledgements

This work has been supported by the National Institute of Arthritis and Musculoskeletal and Skin Diseases (NIAMS) and the National Eye Institute (NEI) of the National Institutes of Health (NIH) under award numbers R01AR072621 and R21EY029081 (PJK and MIK). The content is solely the responsibility of the authors and does not necessarily represent the official views of the National Institutes of Health. This work was also supported by the National Foundation for Ectodermal Dysplasias (NFED; PJK and MIK). We would like to thank Dr. Munkhtuya Tumurkhuu and Neha Joseph (East Carolina University) for conducting preliminary migration assays. We would also like to thank the patients who supported this research by providing skin biopsies.

## Author contributions

PJK and MIK conceptualized and designed this work; DKG and GML obtained skin biopsies; MIK and DKG obtained regulatory approval; SW and MNS generated iPSC-K; JAG, CES, and MNS performed RNA and protein analyses; CES and MNS performed adhesion and migration studies, MNS and SPP performed immunostaining; SW generated the animals; all authors have read and approved the final version of the manuscript.

