## Supplemental Methods for "Effects of TP63 Mutations on Keratinocyte Adhesion and Migration"

**Supplemental Data - METHODS**

**Generation of Human Induced Pluripotent Stem Cell (iPSC)-Derived Keratinocytes (iPSC-K)**

We obtained skin biopsies from AEC patients carrying *TP63* mutations. Biopsies were obtained under IRB protocols #17240 through St Louis University (approval date 6/7/2011), #19-0501 through the University of Colorado Denver (approval date 4/19/2019), and #201901088 through Washington University (approval date 5/13/2019). All research subjects provided their written informed consent/assent prior to their inclusion in the study. We isolated skin fibroblasts from these biopsies and generated iPSC using an integration-free Sendai reprogramming system (Dinella et al., 2018). iPSC lines were tested for their ability to generate cells representing all three germ layers [endoderm, mesoderm, ectoderm (STEMdiff, Stemcell Technologies cat # 05230)]. All iPSC used had a normal karyotype and were free of mycoplasma. Using CRISPR/CAS and TALEN technology, we corrected the *TP63* mutations in each iPSC line, thereby generating pairs of cell lines that are identical except for the presence/absence of a *TP63* mutation. Three pairs of AEC and gene-corrected iPSC lines were used in this study. *TP63* mutations in the AEC iPSC lines are F513S, I537T, and R598L. To generate keratinocytes (iPSC-K), we exposed iPSC-derived embryonic bodies to retinoic acid and bone morphogenetic protein 4 (BMP4). A detailed description of the methodology can be found in (Koch et al., 2022). iPSC-K used in the present study expressed proteins present in basal (undifferentiated) human keratinocytes, such as TP63 and keratin 14 (KRT14). Only iPSC-K preparations that were ≥ 95% positive for TP63 and KRT14 were used for further analysis.

**Generation of TP63 Transgenes and Transduction of Mouse Embryonic Stem (ES) Cells**

Lentiviral vectors co-expressing *TP63-AEC* cDNA with the fluorescence protein dTomato were constructed based on the pHIV-TdTomato plasmid (Addgene cat# 21374). In our constructs, a CAG promoter drives expression of a *TP63-AEC* cDNA and the TdTomato reporter. A T2A self-cleaving peptide sequence was inserted between the coding sequences of *TP63* and TdTomato, resulting in the presence of equimolar amounts of both proteins in transduced cells. Lentiviral stocks were generated using these constructs by SignaGen laboratories.

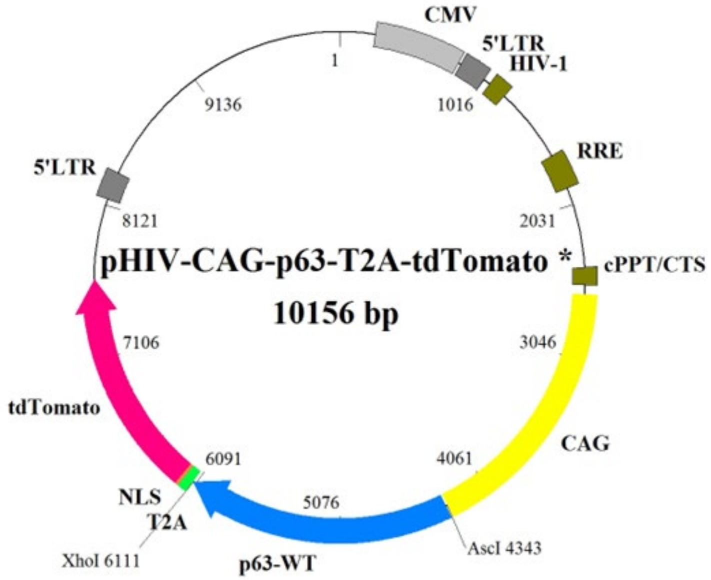


Schematic representation of the lentiviral construct generated to co-express *TP63* with the fluorescent protein TdTomato. Lentiviral vector construction was done in collaboration with the Transgenic Vector Core at the University of Colorado Denver (Dr. Wallace Chick, Core Director).

**Generation of Chimeric Mice Expressing an AEC-Variant of TP63**

129Sv/J derived mouse embryonic stem (ES) cells were transduced in suspension with lentiviral constructs at a MOI (multiplicity of infection) of 50. 72 hours post transduction, ES cells were subjected to FACS sorting using TdTomato expression as a marker for transduced cells. FACS sorting was performed by the Flow Cytometry Core at East Carolina University. Transgenic (TdTomato$^{+}$) ES cells were injected into C57Bl6/N-derived blastocysts, which were then implanted into pseudo pregnant CD-1 female mice (Charles River). Isolation of blastocysts, ES cell injections, generation of pseudo pregnant recipient mice and surgical procedures were done essentially as described (Behringer et al., 2014). Chimeric mice derived using this procedure were identified by PCR detection of the TdTomato transgene in genomic DNA isolated from mouse tail. The following primers were used to detect the TdTomato transgene [forward: 5'-TCCCCGATTACAAGAAGCTG; reverse: 5'-TGGCCATGTAGATGGTCTTG]. Transgenic cells were identified *in vivo* by staining tissue sections with a TdTomato antibody (Origene cat #AB8181-200) (see also below). All animal procedures were approved by the Institutional Animal Care and Use Committee (IACUC) of East Carolina University.

**qRT PCR**

iPSC-K were cultured in CNT-07 media (CellnTec). RNA was isolated using the RNeasy® Plus Mini Kit (Qiagen, cat #74134). cDNA was generated using the iScript cDNA synthesis kit (Bio-Rad, cat #1708841). qRT-PCR was conducted using SsoAdvanced Universal SYBR® Green Supermix (Bio-Rad, cat #1725271), using a CFX Connect Real-Time System (Bio-Rad). Gene expression was normalized to GAPDH and the fold change in expression was calculated using the ddCt method.

Primers used for qRT-PCR

| Gene | Forward (5' to 3') | Reverse (5' to 3') |
|---|---|---|
| ITGA2 | TTAGCGCTCAGTCAAGGCAT | CGGTTCTCAGGAAAGCCACT |
| ITGA3 | CCGGTGCCTACAACTGGAAAG | TGCCTACCTGCATCGTGTACC |
| ITGA6 | TGTGCTTGCTCTACCTGTCG | ACGAGCAACAGCCGCTT |
| ITGB4 | GCCTACGAGGTCTGCTATGG | AGCAGCATCCGGTTCTTAGG |
| COL17A1 | TCACGTTACCCGCCATGC | GTGTTTGACTCCGTCCTCTGG |
| DST | ACAGAAGATGCTGGTGTCCG | TTCTTCGGCGTTTCACTGGA |
| PLEC | AAAGAGAACCAGCTCGGAGG | CGCGGAGGTCTTCATACAGG |
| LAMA3 | AGTTCACAGCAAAGGGT | ACCGTCCGGTATACAAGCCT |
| LAMB3 | CCCATGAATGCCAAAGGTGC | CTGACACCGCTCACAGTTCT |
| LAMC2 | AGCCAAGAACGCTGGGGTTA | AGACCAGCCCCTCTTCATCTA |
| GAPDH | AGCCACATCGCTCAGACAC | GCCCAATACGACCAAATCC |

**Western Blotting**

Western blots were performed using total cell lysates of iPSC-K cultured in CNT-07 media. Briefly, after washing the cells with PBS, cells were lysed in 2x Laemmli Buffer supplemented with 100mM DTT. Primary antibodies used were ITGA6 (Millipore, MAB1982, 1:500), ITGB4 (Cell Signaling, 4707, 1:1000), COL17A1 (Courtesy of Dr. Hendri Pas, University of Groningen, The Netherlands), and GAPDH (Cell Signaling Technology, 5174S). Secondary antibodies (IRDye® 800CW and IRDye® 680RD) were purchased from LiCor, and imaging was performed with the LiCor Odyssey FC Imaging System. The Empiria Studio software suite (Li-Cor) was used for protein visualization and quantification.

**Plating Efficiency Assay**

96-well plates (Falcon, cat# 353072) were coated with extracellular matrix (ECM) proteins: laminin 332 (BioLamina, LN332-0502), laminin 511 (Nippi Matrixome, iMatrix-511), or collagen 4 (Sigma-Aldrich, C5533-5MG) for either 1 hour at room temperature or overnight at 4°C. iPSC-K were labeled with a cell tracking dye (Abcam, ab138891) for 30 minutes following manufacturers' recommendations. After dissociation with Accutase (Stemcell Technologies, cat # 07920), 20,000 cells were plated in each ECM-coated well. After 30 minutes, the wells were washed with PBS and attached cells were quantified either based on fluorescence (Fluoroskan, Ascent; Thermo Scientific) or by directly counting cell numbers using Image J (NIH). Two-tailed unpaired Student t-tests were performed to determine statistical significance. $p<0.05$ was considered statistically significant.

**Migration**

A 96-well plate-based cell migration assay from Platypus Technology was used (CMA5.101). The wells of the culture dish were coated with extracellular matrix proteins as described above. Next, Oris™ Cell Seeding Stoppers were inserted into the wells, thereby creating a circular area to which cells cannot attach in the following steps. iPSC-K were labeled with a cell tracking dye (Abcam; ab138891) for 30 minutes. The cells were then washed with PBS followed by dissociation with Accutase. 30,000 cells were seeded in each well of the 96-well plate containing the Seeding Stoppers. After overnight culture, the Seeding Stoppers were removed to create an ECM-coated area onto which the cells can migrate. Cells were imaged at various time points with a Nikon Eclipse TS2 microscope, and migrating cells were quantified using the Image J software package. Two-tailed unpaired Student t-tests were performed to determine statistical significance. $p<0.05$ was considered statistically significant.

**Immunofluorescence Microscopy**

Mouse tissues were harvested on embryonic day 18.5, formalin-fixed, and paraffin-embedded. Tissue sections were generated by the Brody School of Medicine Histology Core. Immunofluorescent staining was performed following standard procedures (Koster et al., 2009). Immunofluorescence on mouse skin was performed with primary antibodies against ITGA2 (Abcam, ab133557, 1:500) and TdTomato (see above). AEC patient skin samples were stained with primary antibodies against ITGA2 and COL17A1. Secondary Alexa Fluorochrome-labeled antibodies were purchased from Invitrogen. Slides were mounted with DAPI Fluoromount-G® (SouthernBiotech, 0100-20) and imaged on a Nikon Eclipse Ni microscope using the NIS-Elements BR 2.21.02 imaging software.
